# Multi-scale histology reveals localized ageing signatures in normal human colon

**DOI:** 10.64898/2026.10.20.760359

**Authors:** Gilad Altshuler, Coral Teper, Yoash Barak, Sarit Aviel-Ronen, Alex Loebel, Iris Barshack

## Abstract

Ageing is accompanied by heterogeneous changes in tissue structure, cellular composition, and microenvironments, yet how these changes are reflected in routine human histology remains poorly understood. Here, we introduce a histology-first, multi-scale digital pathology framework to study human colon ageing using hematoxylin and eosin (H&E) whole-slide images from clinically normal colon biopsies. The framework combines pathology foundation-model patch embeddings, attention-based multiple-instance learning, cell-level morphometric profiling, and prototype-based tissue compartmentalization. We show that chronological age can be predicted from normal colon tissue using mesoscopic representations derived from pathology foundation-model patch embeddings and attention-based multiple-instance learning. Incorporating cell-type-level morphometric features further improves prediction and refines the spatial allocation of model attention. By integrating attention maps with prototype-defined tissue compartments, we identify localized age-associated differences within stromal, epithelial, and lymphoid mucosal niches. Finally, we show that prediction-discordant cases exhibit cellular profiles shifted toward the model-assigned age group, suggesting that some apparent model errors may reflect tissue-level age discordance beyond chronological age. Together, these findings show that routine H&E histology captures spatially localized signatures of normal human colon ageing and provides a scalable framework for linking tissue morphology with biological ageing state.

## 1 Introduction

Ageing is accompanied by progressive molecular, cellular, and tissue-level changes that reduce physiological resilience and increase susceptibility to disease. Canonical frameworks describe ageing through interconnected hallmarks, including genomic instability, epigenetic alterations, cellular senescence, stem-cell exhaustion, altered intercellular communication, chronic inflammation, and dysbiosis [1, 2]. However, ageing is not fully captured by chronological time. Individuals of the same chronological age can differ markedly in biological state, and increasing evidence suggests that biological ageing is both tissue-specific and spatially heterogeneous, with different organs and local microenvironments ageing at distinct rates [3]. This has motivated the development of ageing clocks and tissue-specific measures of biological age [4, 5]. While molecular clocks have provided important insights, less is known about how ageing is encoded in routine tissue morphology, particularly in histologically normal human tissues.

The colon is an especially relevant organ for studying tissue ageing [6–8]. It is organized into spatially distinct compartments, including crypt epithelium, lamina propria, stromal regions, lymphoid structures, and deeper submucosal tissue. The colonic epithelium undergoes continuous renewal and contributes to barrier maintenance, while the mucosa contains a dense immune network that interacts with the microbiome and luminal environment. Ageing has been associated with reduced regenerative capacity of intestinal stem cells, altered epithelial turnover, barrier dysfunction, stromal remodeling, immune-cell redistribution, and chronic low-grade inflammatory activation (“inflammaging”) [7, 9–11]. Recent spatial and single-cell studies further suggest that colon ageing is regionally structured, reflecting coordinated changes across epithelial, stromal, vascular, immune, and stem-cell niches rather than uniform tissue-wide decline [12, 13]. These observations make the colon a compelling model for studying how ageing manifests across tissue architecture, cellular composition, and local microenvironments.

At the same time, computational pathology has reached a critical point. Large pathology foundation models can now extract powerful representations from H&E whole-slide images (WSIs) and support diverse downstream tasks across cancer diagnosis, prognosis, biomarker prediction, and tissue phenotyping [14–17]. Recent work has begun to extend these approaches beyond neoplastic pathology. For example, Meng et al. showed that machine-learning models trained on histological images from deceased donors can predict chronological age and identify image-associated genetic loci [18]. Daly et al. provided a complementary high-resolution view of mice colon ageing using spatial transcriptomics and single-nucleus RNA sequencing, revealing spatiotemporal cellular and molecular programs across the ageing colon [13]. In parallel, Peng et al. benchmarked pathology foundation models in non-neoplastic placental pathology, highlighting the promise of foundation-model representations beyond cancer-focused applications [19]. Together, these studies show that histology, spatial molecular profiling, and foundation models can capture biologically meaningful variation in non-neoplastic tissues.

A remaining challenge is to move from slide-level prediction to spatially and biologically interpretable tissue analysis. Attention-based multiple instance learning provides a natural framework for linking slide-level predictions to patch-level relevance maps [20, 21]. Pathology foundation models are now being applied beyond diagnostic classification, serving as representation-learning systems that can capture latent biological structure from routine histology [14, 22–25]. Prototype-based approaches such as PANTHER allow histological patches to be grouped into recurring morphological patterns or tissue compartments, enabling interpretation of model behavior in relation to tissue architecture [26]. In parallel, cell-detection and classification methods such as ClassPose provide explicit measurements of cell-type abundance and morphology from H&E images [27]. Together, these developments create an opportunity to combine mesoscopic tissue representations with microscopic cellular information for interpretable ageing analysis.

Despite this opportunity, existing histology-based ageing studies have primarily focused on prediction or genetic association, whereas spatial omics studies provide deep molecular resolution but are costly and difficult to scale across large clinical cohorts. Moreover, to our knowledge, no prior study has integrated foundation-model histology embeddings, cell-level morphology, attention-based localization, and compartment-aware analysis to study biological ageing in normal human tissue.

Here, we address this gap by developing a histology-first, multi-scale framework for studying human colon ageing from routine H&E biopsy slides. The approach integrates attention-based multiple instance learning, foundation-model patch embeddings, ClassPose-derived cellular and morphometric features, and PANTHER-based tissue compartmentalization [26, 27]. We first test whether chronological age can be predicted from histologically normal colon tissue, then ask whether adding cell-level information improves prediction and changes where the model attends. Finally, we use attention-guided, compartment-aware analysis to identify age-associated cellular and morphometric differences within stromal, epithelial, and lymphoid regions. By linking slide-level prediction to localized cellular and compartment-specific features, this work provides a foundation for scalable histology-derived biomarkers of biological tissue ageing and age-related mucosal dysfunction.

## 2 Results

### Cohort assembly and study design

We assembled an age-spanning cohort of patients who underwent colon biopsy as part of routine clinical care. Biopsy specimens were processed using standard histopathology workflows, including H&E staining, whole-slide digitization, and extraction of associated clinical metadata (Fig. 1a). To focus the analysis on baseline ageing-related morphology, we applied inclusion and exclusion criteria to retain only specimens classified as normal colon and to exclude samples with abnormal pathology or potentially confounding disease processes. Slides were further filtered at the image level to remove empty, blurred, non-viable, or incorrectly sampled tissue.

**Fig. 1.**
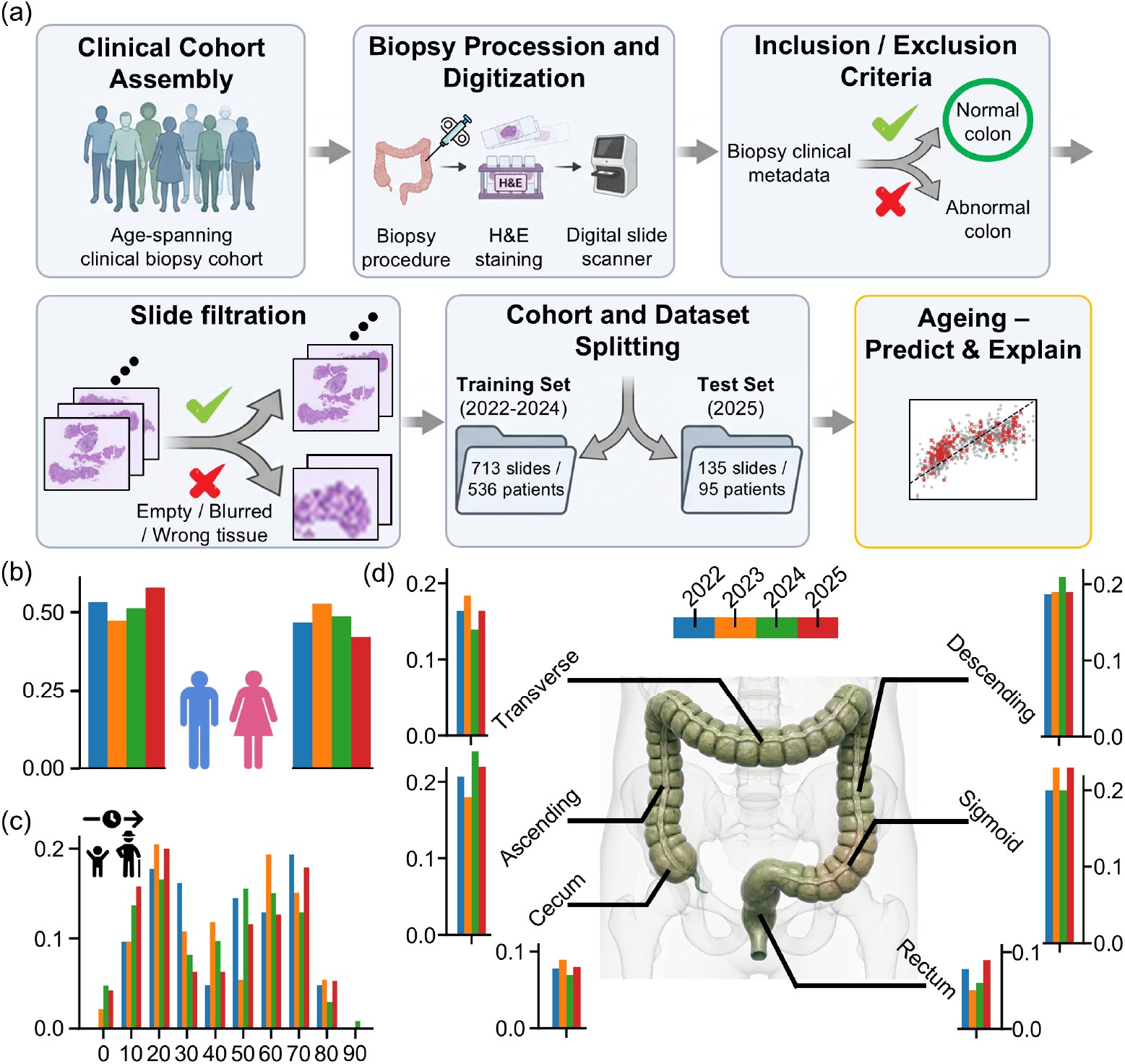
Cohort and study design overview. (a) Overview of cohort assembly and data curation. An age-spanning clinical biopsy cohort was assembled from patients undergoing colon biopsy, followed by H&E staining, slide digitization, metadata extraction, inclusion/exclusion filtering for normal colon tissue, slide-level quality control, temporal train/test splitting, and downstream ageing prediction and interpretation. Slides from 2022-2024 were used for training, and slides from 2025 were held out for testing. (b) Case-level sex distribution across acquisition years. (c) Case-level age distribution across decade-level age bins and acquisition years. (d) Distribution of biopsy location across colon anatomical regions and acquisition years.

To obtain a temporally separated evaluation design, slides from 2022-2024 were used for training, whereas patients and slides from 2025 were held out as an independent test set. After filtering, the training set contained 713 slides from 536 patients, and the test set contained 135 slides from 95 patients (Fig. 1a). This split was used for all downstream model development, evaluation, and interpretability analyses.

We next examined key cohort characteristics across acquisition years. Sex distributions were broadly balanced across years (Fig. 1b), and the cohort covered a wide age range with representation across decade-level age bins (Fig. 1c). Biopsy locations spanned the major anatomical regions of the colon, including cecum, ascending, transverse, descending, sigmoid, and rectum, with similar location distributions across years (Fig. 1d). Together, this design enabled assessment of age-related histological variation while maintaining a held-out temporal test set for model evaluation.

### Multi-scale modeling of colon ageing

To capture age-associated morphological variation across levels of tissue organization, we developed a multi-scale framework that integrates mesoscopic tissue architecture with microscopic cellular features (Fig.2). The pipeline is organized into three stages-feature extraction, model training, and interpretation-enabling both predictive modeling and spatially resolved biological analysis.

**Fig. 2.**
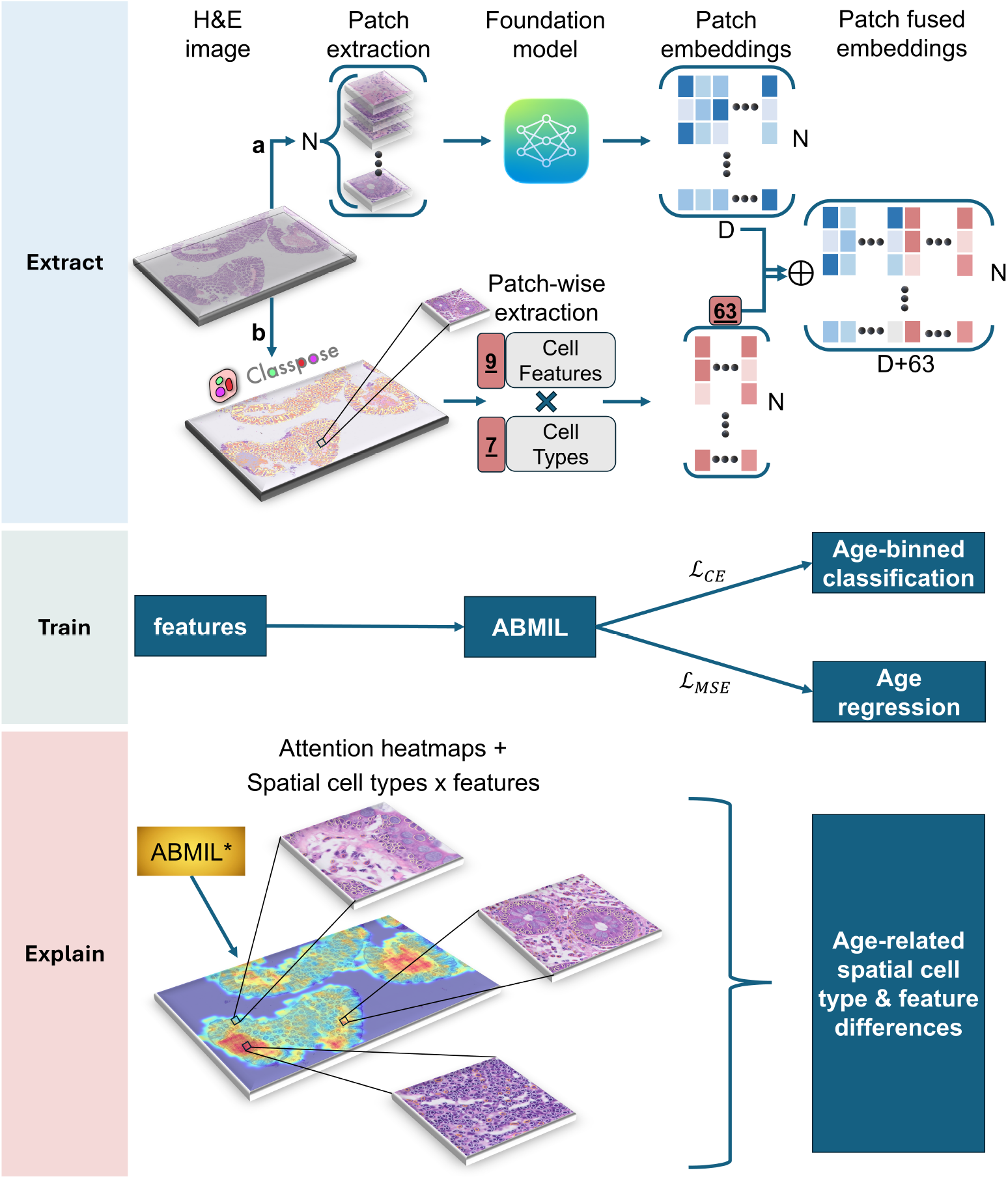
Multi-scale fusion framework for colon ageing analysis. Whole-slide H&E images are analyzed using complementary mesoscopic and microscopic representations that are integrated through a fusion model to enable prediction and biological interpretation. (a) WSIs are partitioned into patches, which are embedded using a pretrained foundation model to generate patch-level feature representations. (b) In parallel, cellular composition and morphometric features are extracted using ClassPose [27], yielding per-patch descriptors across multiple cell types and features. These repre-sentations are combined via feature-level fusion, producing enriched patch embeddings that integrate contextual tissue architecture with cellular information. The fused features are aggregated using an attention-based multiple instance learning (ABMIL) model to perform both age-binned classification and continuous age regression. For interpretability, attention weights are projected back onto the tissue to generate spatial heatmaps, which are jointly analyzed with cell-type distributions and morphometric features. This enables localization of age-associated remodeling to specific histological regions and cell populations, including epithelial, immune, and stromal compartments.

Whole-slide H&E images were first tessellated into non-overlapping patches, which were embedded using a pretrained foundation model (e.g., UNI [14] or Virchow [15, 16]) to obtain high-dimensional representations capturing mesoscopic tissue context. In parallel, we extract microscopic cellular features using ClassPose [27], which performs cell detection and classification directly on the tissue.

For each patch, the cellular composition and morphology across seven cell categories (six annotated cell types and an aggregate “total” category) were quantified. Within each category, we derived nine features describing cell density and morphology, including measures of abundance, area, and shape (e.g., circularity, elongation, and perimeter-to-area ratio). This resulted in a compact set of 63 cell-derived features per patch, capturing local cellular organization and structural variation.

The mesoscopic embeddings and microscopic cellular features were then combined via feature-level fusion to generate enriched patch representations that jointly encode tissue architecture and cell-level information (Fig. 2).

Fused patch-level features were aggregated into a slide level representation using an attention-based multiple instance learning (ABMIL) framework [20]. This approach enables the model to assign differential importance to individual patches while producing a global prediction for each slide.

The model was trained to predict age using both classification (age-binned) and regression (continuous age) objectives. This dual formulation allows the model to capture coarse-grained age-related patterns while using regression to validate the strength and continuity of the underlying signal. Importantly, the attention mechanism provides a natural means to identify spatial regions that contribute most strongly to the prediction.

To interpret the learned representations, attention weights were projected back onto the tissue to generate spatial heatmaps highlighting informative regions. These regions were then analyzed in conjunction with the underlying cellular features, enabling joint characterization of *where* age-related signals are localized and *what* cellular and morphometric patterns drive them.

This integrated framework allows us to link predictive performance with biologically interpretable features, facilitating the identification of age-associated remodeling across epithelial, immune, and stromal compartments.

### Multi-scale fusion improves predictive performance and reveals structured age-related errors

The contribution of mesoscopic and microscopic features were subsequently evaluated through a series of experiments (Fig. 3a). Models based on attention-derived patch embeddings achieved strong performance (AUC = 0.895), while incorporation of cellular features extracted by ClassPose further improved performance (AUC = 0.932), demonstrating complementary information between tissue architecture and cell-level descriptors. In contrast, models relying solely on cellular features or non-attentive aggregation (e.g., mean pooling) showed significantly reduced performance, highlighting the importance of both spatial context and learned patch weighting.

**Fig. 3.**
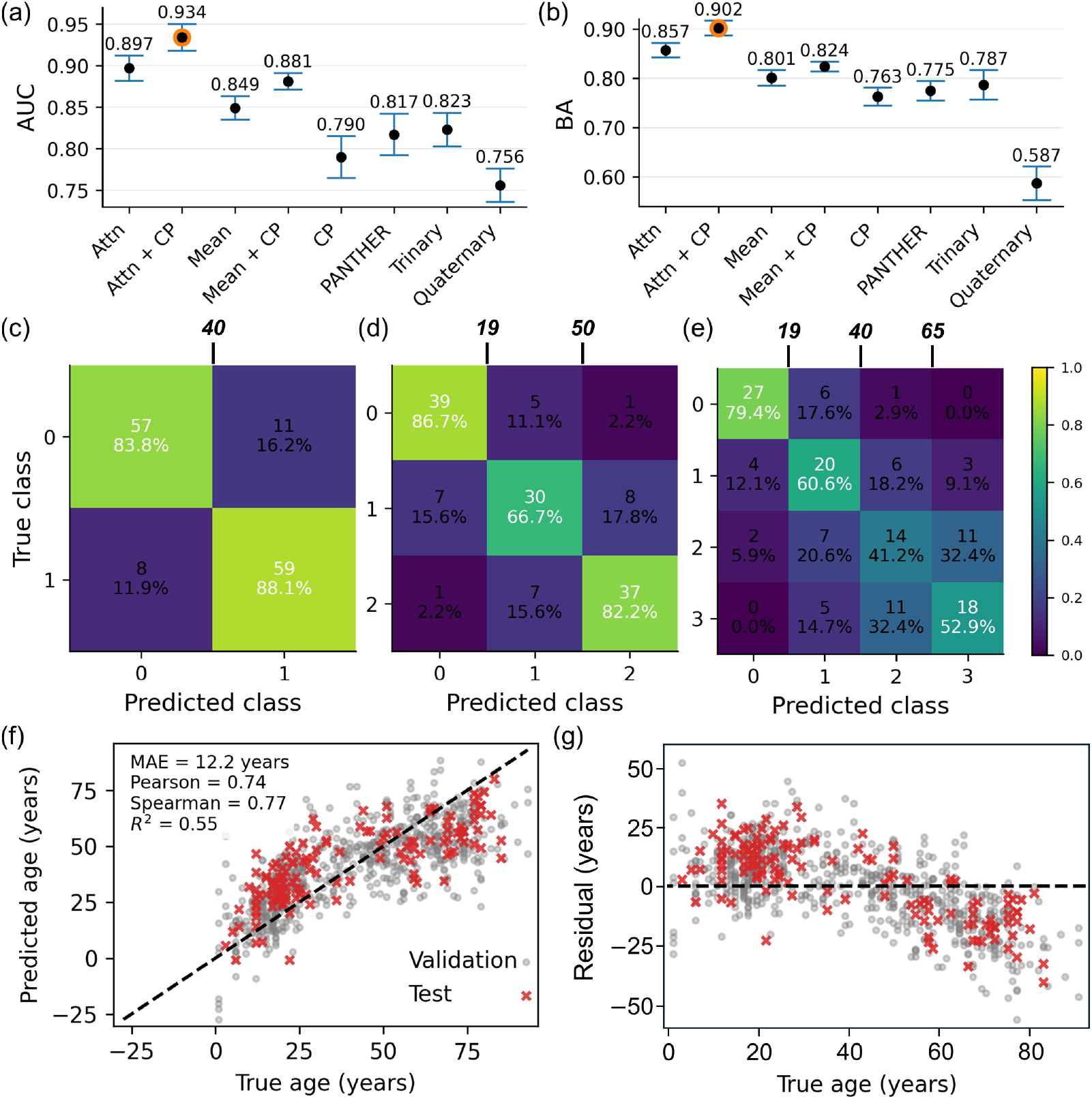
Model performance and ablation analysis. (a,b) Performance comparison across model variants using (a) area under the receiver-operating characteristic curve (AUC) and (b) balanced accuracy (BA). Attn denotes attention-based multiple instance learning (ABMIL) applied to patch embeddings, while CP denotes cellular features extracted using ClassPose. The proposed fusion model (Attn + CP) combines both representations. Baselines include attention-only (Attn), mean pooling (Mean) as a non-attentive aggregation alternative, cellular features alone (CP), their corresponding combinations, and PANTHER aggregation and classification [26].Trinary and quaternary classification settings are also shown. Markers denote the mean performance across *N* = 10000 test-set bootstrap runs, and error bars indicate the standard deviation. (c-e) Confusion matrices for (c) binary, trinary, and (e) quaternary age classification. Cells display counts and percentages. Age thresholds used for binning are indicated above each matrix. (f) Regression performance of the ABMIL model predicting continuous age, with performance metrics shown. (g) Residuals of the regression model as a function of true age.

This trend was consistent across evaluation metrics, including balanced accuracy (Fig. 3b), and extended to multi-class settings. Increasing the granularity of age prediction from binary to trinary and quaternary classification led to a gradual reduction in performance, reflecting the increased difficulty of finer age stratification. Notably, however, the relative performance ranking between models was preserved, with the fused model consistently outperforming baselines.

Analysis of confusion matrices revealed a structured pattern of misclassification (Fig. 3c-e). Errors were predominantly confined to adjacent age groups, with a marked decrease in predictions assigned to distant classes. For example, in the quaternary setting, samples from the oldest age group were most frequently misclassified as the neighboring group, and rarely assigned to younger categories. This pattern indicates that the learned representations capture a continuous ageing signal rather than discrete class boundaries.

To further assess this continuity, we evaluated the model in a regression setting (Fig. 3f). Predicted age showed strong correlation with chronological age (Pearson *r* = 0.74, Spearman *r* = 0.77, *R*^2^ = 0.55, MAE = 12.2 years), supporting the presence of a smooth underlying relationship. Residual analysis (Fig. 3g) revealed no abrupt discontinuities, with errors varying gradually across the age range. Together, these results suggest that while classification provides a convenient framework for interpretability, the model captures a continuous, biologically meaningful ageing trajectory.

### Mesoscopic tissue composition is stable across age despite biologically meaningful compartmentalization

To investigate whether age-associated differences are reflected at the level of mesoscopic tissue organization, we analyzed patch-level prototype assignments using PANTHER [26]. This approach partitions histological patches into a discrete set of recurring tissue phenotypes, enabling a coarse-grained description of tissue composition across WSIs.

Visual inspection of prototypical assignment maps revealed that the learned proto-types correspond to biologically meaningful tissue structures (Fig. 4a). These included distinct compartments such as epithelial crypts, connective tissue, lymphoid aggregates, and submucosal regions, as illustrated by representative image patches for each prototype (Fig. 4c). At the local level, regions of interest exhibited coherent proto-type assignments and characteristic prototype distributions, indicating that the model captures spatially consistent histological patterns.

**Fig. 4.**
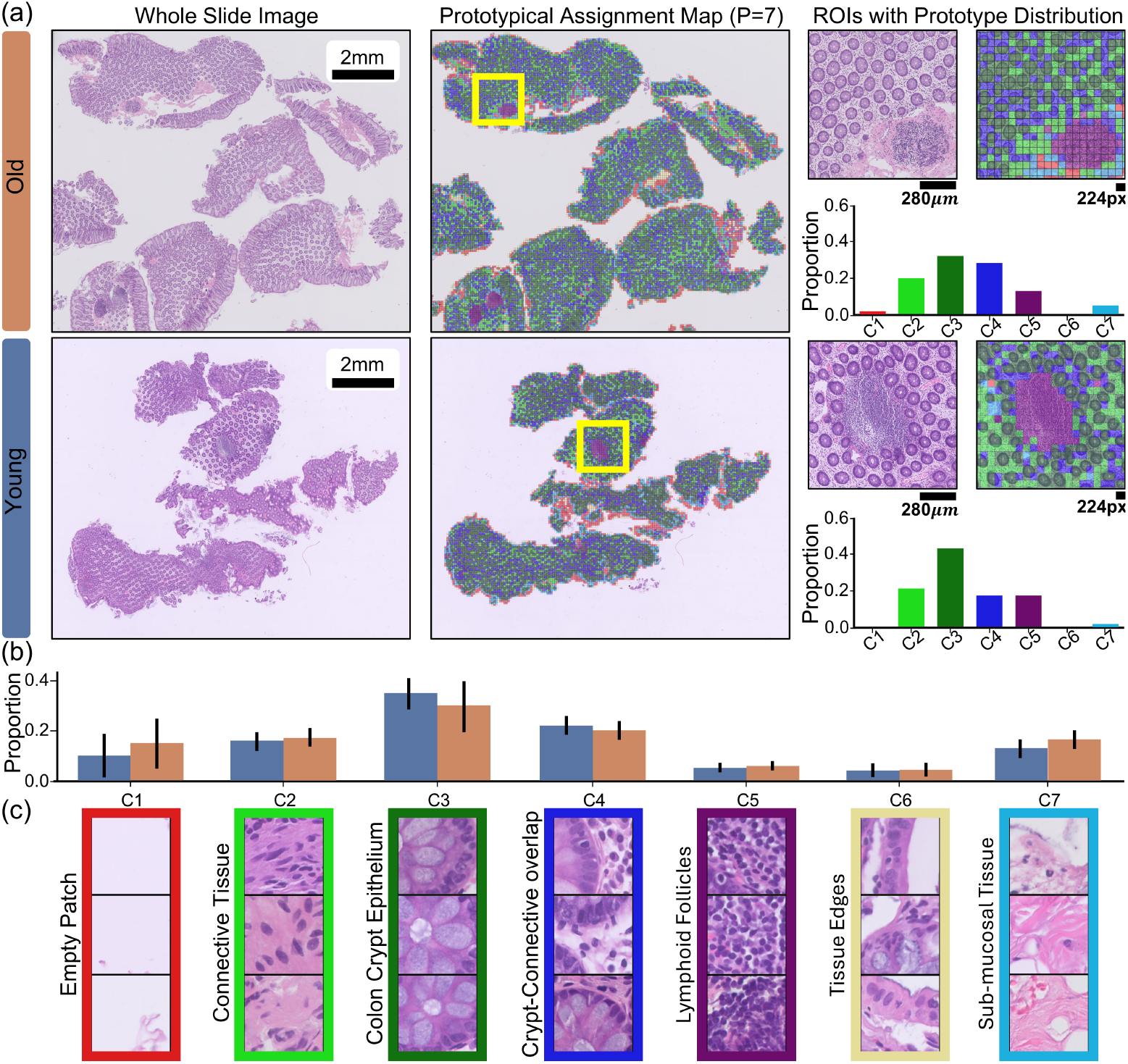
Prototype-based interpretation of tissue organization. (a) Representative WSIs from an old (67 years old) and a young (20 years old) individuals, alongside their corresponding PANTHER prototypical assignment maps (*C* = 7). Each patch is colored according to its assigned prototype. Highlighted regions of interest (ROIs) are shown with higher magnification, together with their local prototype assignment maps and corresponding prototype proportion histograms. (b) Distribution of prototype proportions across the test cohort, stratified by age group (young vs. old). Bars represent mean proportions across slides, with error bars indicating standard deviation. No significant differences are observed between age groups. (c) Representative image patches corresponding to each prototype (C1-C7), with manually assigned biological interpretations.

Despite this interpretability, we observed minimal differences in the global distribution of prototype assignments between young and old cohorts. Across the test set, prototype proportions were largely conserved between age groups (Fig. 4b), with no substantial shifts in the relative abundance of any individual prototype.

These findings suggest that, while mesoscopic tissue compartmentalization captures meaningful histological structure, it alone is insufficient to explain age-associated variation in colon tissue. This is consistent with the improved predictive performance observed when incorporating higher-resolution cellular features (Fig. 3a,b), indicating that age-related signals are likely encoded at finer spatial scales beyond coarse tissue composition.

### Fusion redistributes attention toward spatially localized histological regions

To examine how the integration of cell-level features alters model interpretability, we compared the spatial distribution of attention weights between the attention-only model and the fused model incorporating ClassPose-derived cellular features. Visual inspection of representative whole-slide heatmaps show that the fused model produce sharper and more spatially localized attention patterns, with increased emphasis on discrete tissue regions rather than broadly distributed activation (Fig. 5a,b).

**Fig. 5.**
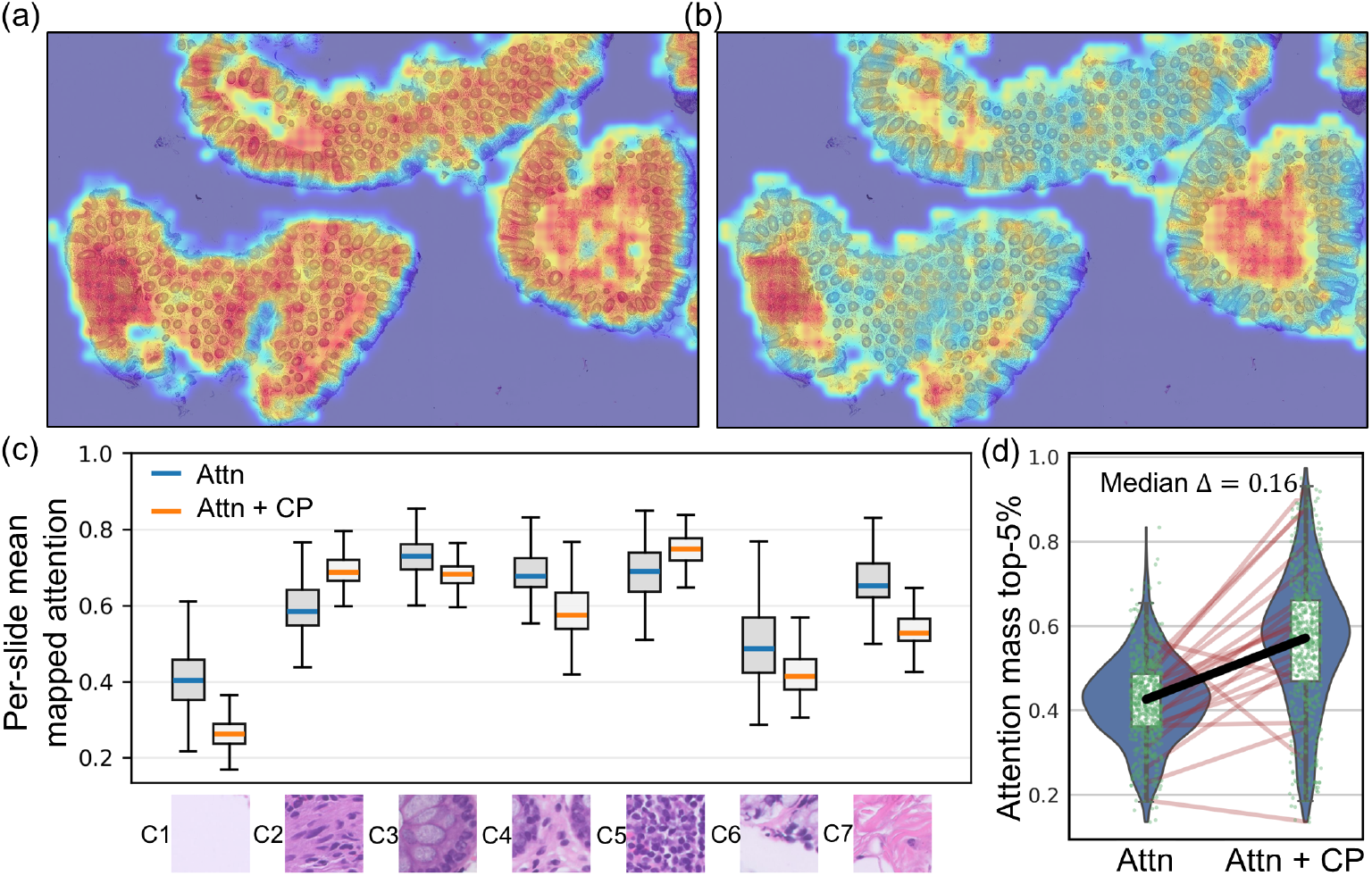
Integration of cell-level features concentrates attention into spatially localized regions. (a) Representative whole-slide attention heatmaps from the attention-only model and (b) the fused model incorporating ClassPose-derived cellular features. The fused model exhibits more spatially localized and sharper attention patterns. (c) Per-slide mean mapped attention values across histological features for both models, illustrating systematic shifts in attention allocation following fusion. (d) Distribution of the fraction of total attention mass assigned to the top 5% of patches per slide. Each point represents a slide, with paired lines connecting the same slide across models. The fused model assigns significantly more attention mass to top-ranked patches (median Δ= 0.16).

To characterize where this redistribution occurs, the mean mapped attention was next summarized across histological feature categories, extracted with PANTHER [26]. This analysis revealed systematic shifts in attention allocation following fusion, indicating that the addition of cellular information changes not only predictive performance, but also the types of tissue regions prioritized by the model (Fig. 5c).

Attention concentration was quantified by computing, for each slide, the fraction of total attention assigned to the top-ranked patches (by attention weights). Across slides, the fused model consistently allocated a larger share of attention to these top patches compared to the attention-only model (Fig. 5d). For example, the median fraction of attention mass captured by the top 5% of patches increased after fusion, indicating that a smaller subset of patches accounts for a larger portion of the model’s decision. Similar results were observed in the top 1% and 10% patches analysis (Supplementary Fig. 2a,b)

As a complementary summary of the same effect, normalized attention entropy decreased in the fused model (Supplementary Fig. 2c), consistent with a more peaked attention distribution. Together, these results show that integrating cell-level features sharpens the spatial allocation of model attention and concentrates decision weight into a more localized subset of histological regions.

### High-attended patches across PANTHER compartments reveal age-associated cellular and morphometric differences

To determine whether model-attended regions contain age-associated cellular differences, we analyzed ClassPose-derived features within PANTHER-defined tissue compartments. For each slide and each PANTHER prototype label, patches were ranked by their mapped attention scores and divided into high- and low-attention subsets using the top and bottom 5% of patches, respectively. Within each slide, prototype label, and attention stratum, ClassPose features were averaged across selected patches, yielding one value per slide for each feature-prototype-attention combination. Age-associated differences were then tested between young and old groups across slides. Feature-prototype combinations were retained for visualization if they satisfied both a statistical threshold (*p <* 0.01) and a minimum effect-size threshold (|Δ| *>* 0.2 in standardized units). This analysis identified significant differences in high-attention patches within three main PANTHER compartments: connective tissue, colon crypt epithelium, and lymphoid follicles (Fig. 6, top).

**Fig. 6.**
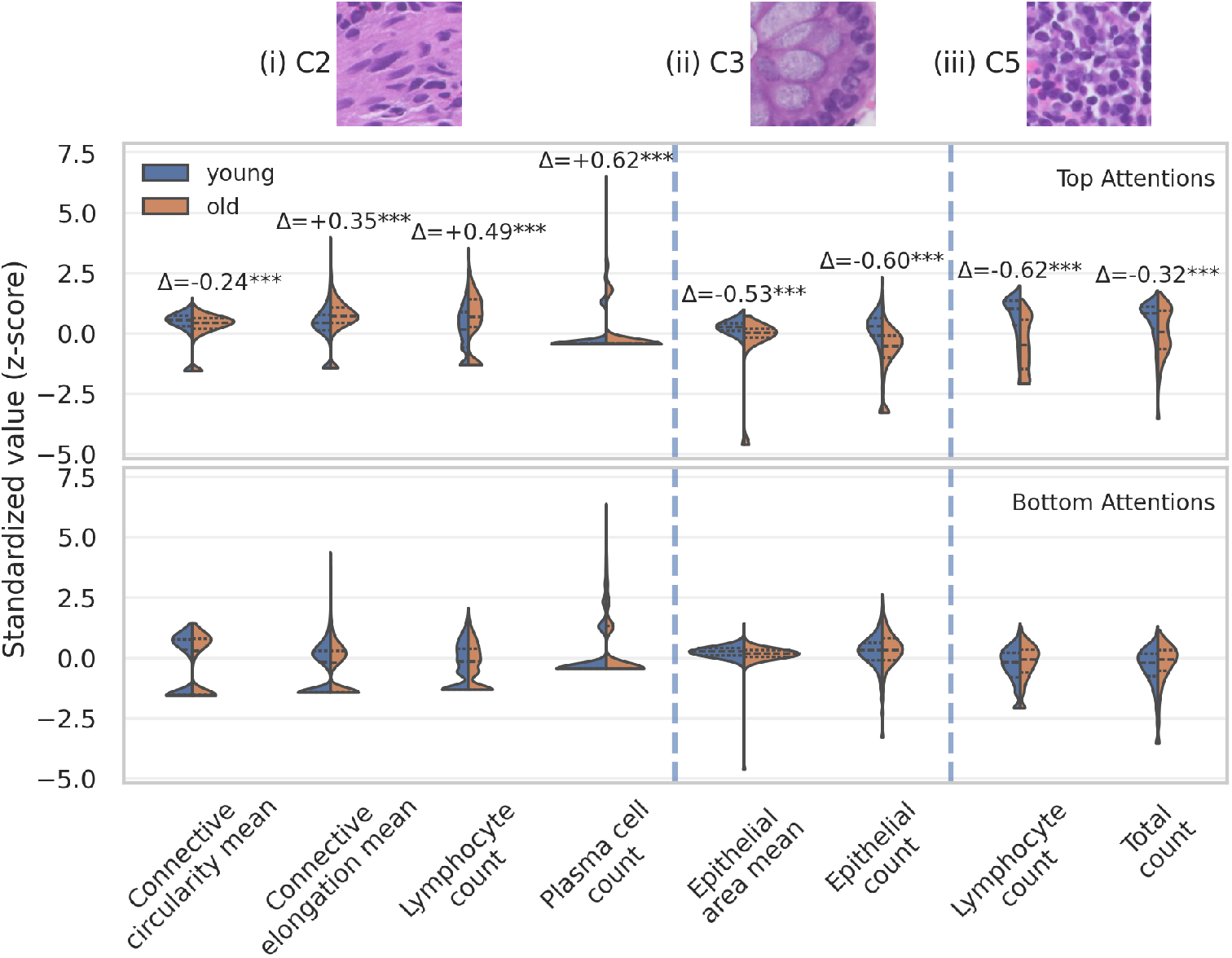
Age-associated cellular and morphometric differences are enriched in high-attention tissue compartments. ClassPose-derived features were computed at the patch level and analyzed within PANTHER-defined tissue compartments. For each slide and each prototype label, patches were stratified into high- and low-attention subsets using the top and bottom 5% of mapped attention scores, respectively. Feature values were averaged per slide within each prototype label and attention stratum. Comparisons between young (blue) and old (orange) cohorts were performed across slides for each feature-prototype combination. Only combinations satisfying both statistical significance (*p <* 0.01) and a minimum effect size (|Δ| *>* 0.2) are shown. **Top**: High-attention patches show significant age-associated differences within selected PANTHER compartments. Representative prototype regions are shown above: (i) stromal/connective tissue, (ii) colon crypt epithelium, and (iii) mucosal lymphoid follicle. **Bottom**: The same feature-prototype combinations show weaker or absent differences in low-attention patches.

In the connective tissue compartment, older samples showed lower mean circularity of connective cells (Δ= −0.24) and higher mean elongation (Δ= +0.35), together with increased lymphocyte count (Δ= +0.49) and plasma cell count (Δ= +0.62). In the colon crypt epithelium compartment, older samples showed lower epithelial mean area (Δ= −0.53) and lower epithelial cell count (Δ= −0.60). In the lymphoid follicle compartment, older samples showed lower lymphocyte count (Δ= −0.62) and lower total cell count (Δ= −0.32).

In contrast, the same feature-prototype combinations showed weaker or absent differences in low-attention patches (Fig. 6, bottom). These results indicate that the detected age-associated cellular and morphometric differences are enriched in model-attended regions rather than uniformly distributed across all tissue patches.

### Prediction discordance reveals tissue-level signatures of chronological versus biological age

Although chronological age provides a convenient and objective label for supervised learning, it is an imperfect surrogate for biological ageing. Individuals of the same chronological age may differ substantially in tissue maintenance, inflammatory burden, cellular composition, and cumulative exposure history. Accordingly, a model trained to classify chronological age may assign some samples to the opposite age class not because the prediction is biologically uninformative, but because the tissue morphology and cellular state of that individual more closely resemble the opposite chronological group.

To examine this possibility, we stratified the same age-associated cellular features (Fig. 6) according to slide-level prediction outcome. For each significant feature-prototype combination identified in high-attention regions, samples were divided by chronological class and model prediction, yielding true negatives, false positives, false negatives, and true positives, with old age treated as the positive class. We then compared the feature distributions of correctly and incorrectly classified samples within each chronological age group.

Across multiple tissue compartments and cellular features, correctly classified samples closely resembled the overall distribution of their chronological class with no significant changes. In contrast, misclassified samples significantly tended to shift toward the distribution of the opposite chronological class. Young samples predicted as old showed feature profiles that were more similar to old samples, whereas old samples predicted as young showed feature profiles that were more similar to young samples (Fig. 7).

**Fig. 7.**
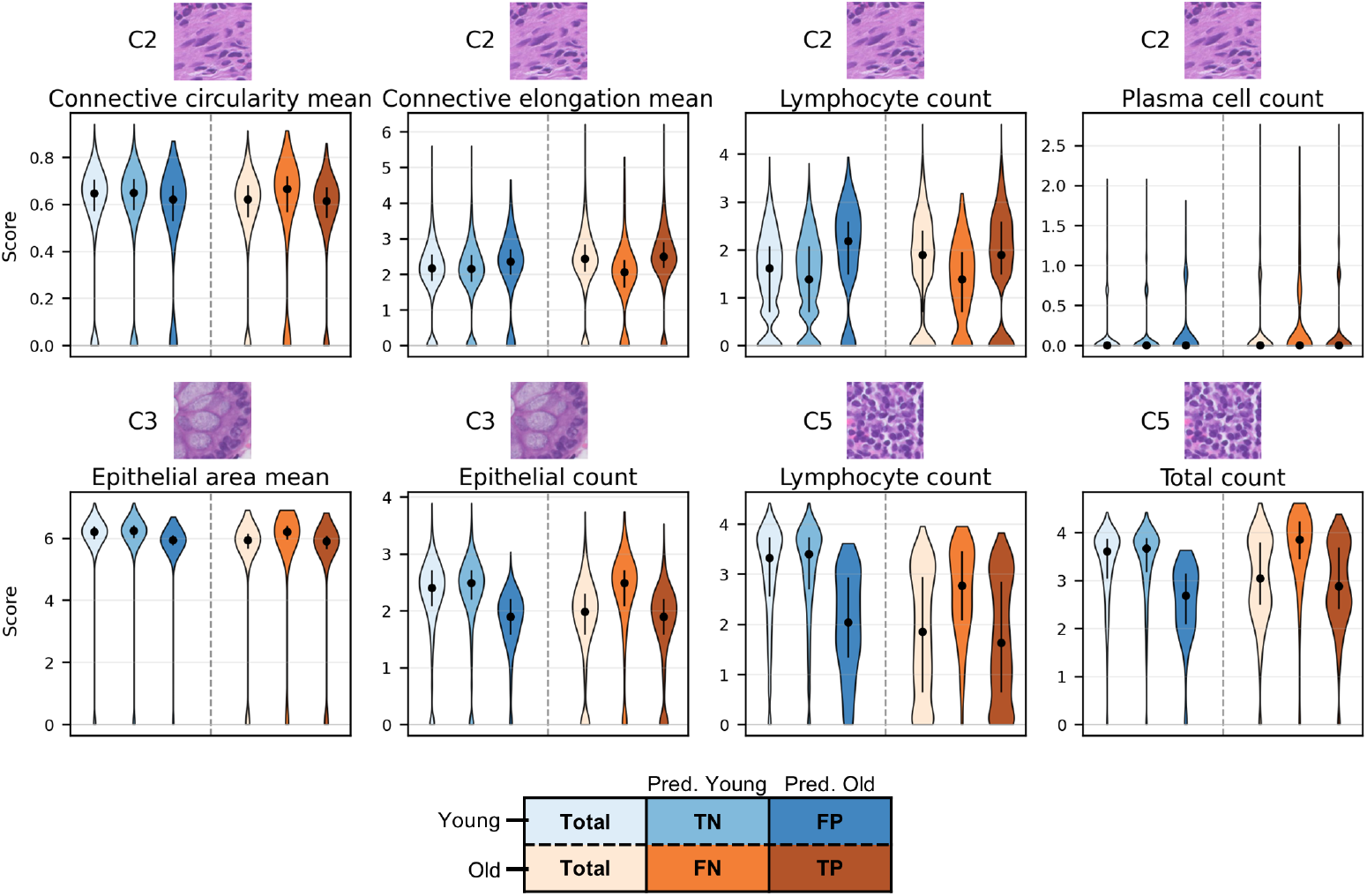
Prediction-outcome stratification of selected ClassPose features in top attention patches. For each significant age-associated feature-prototype combination identified in high-attention regions, feature-score distributions were stratified by chronological age class and model prediction outcome. Chronological young samples are shown on the left side of each panel and chronological old samples on the right side, separated by a dashed vertical line. Within each chronological class, the first violin shows the total class distribution, followed by the distributions of samples predicted as young or old.

These findings suggest that the model’s prediction discordance is not purely random with respect to the underlying histology. Rather, samples that are discordant with their chronological labels show cellular and tissue-compartment features consistent with the model-assigned age class. Because the model was trained using binary chronological age labels, these results do not directly define biological age. However, they support the interpretation that chronological misclassification may partly reflect biological heterogeneity in tissue ageing. In this sense, the discordant predictions provide an internal test of whether the model is capturing tissue-level ageing state rather than only separating chronological age groups.

Together with the spatial enrichment of age-associated features in high-attention regions, this analysis strengthens the link between model attention, cellular remodeling, and biological ageing. It suggests that the learned representation captures a histological ageing axis along which some individuals appear biologically older or younger than their chronological age, consistent with the broader concept of chronological-biological age discordance described in ageing-clock studies.

## 3 Discussion

We present a multi-scale framework for studying human colon ageing from routine H&E whole-slide images. By combining patch-level foundation-model representations, ClassPose-derived cell-level features, attention-based multiple instance learning, and PANTHER-based tissue compartmentalization, our approach links slide-level age prediction with spatially resolved biological interpretation. In particular, we show that cellular features improve predictive performance, refine the spatial allocation of model attention, and help identify age-associated changes within specific stromal, epithelial, and lymphoid tissue compartments. These results suggest that ageing-related morphology in normal colon is not captured solely by global tissue composition, but emerges from localized cellular and morphometric remodeling within defined histological niches.

### Localized ageing-associated remodeling emerges within histological compartments

Rather than treating deep learning predictions as black-box outputs, our framework uses attention-guided and compartment-aware analysis to generate biologically interpretable hypotheses regarding tissue ageing [28, 29].

The cellular and morphometric differences detected in high-attention regions were not randomly distributed across tissue compartments. Instead, they formed a biologically coherent pattern involving stromal, epithelial, and lymphoid mucosal niches. This supports the interpretation that the model is not only predicting chronological age, but also localizing tissue regions in which ageing-related histological variation is enriched.

In connective tissue compartments, older samples showed increased lymphocyte and plasma cell counts together with altered connective-cell morphology, including reduced circularity and increased elongation. The increased lympho-plasmacytic abundance observed within stromal compartments is consistent with the concept of inflammaging, in which chronic low-grade immune activation progressively reshapes tissue microenvironments during ageing. [10, 11, 13, 30–33]. Although our histology-based analysis does not establish fibroblast activation or fibrosis directly, the joint appearance of lymphoplasmacytic enrichment and altered connective-cell shape suggests a plausible stromal-immune ageing phenotype in the lamina propria.

In colon crypt epithelium compartments, older samples showed lower epithelial cell count and reduced epithelial cell area. This is consistent with the known dependence of the intestinal epithelium on continuous renewal, stem-cell activity, and barrier maintenance. Ageing has been linked to altered epithelial homeostasis, impaired barrier function, and changes in mucosal repair programs [6, 13, 34–36]. Thus, the epithelial features identified by the model may reflect local changes in crypt cellularity or epithelial maintenance rather than gross changes in tissue composition.

In lymphoid follicle compartments, older samples showed lower lymphocyte and total cell counts. Gut-associated lymphoid tissues are organized immune structures involved in mucosal antigen sampling and adaptive immune responses [37, 38]. The reduced cellularity observed in lymphoid follicle-like compartments, together with increased lymphocyte and plasma cell abundance in connective tissue compartments, may indicate age-related changes in the organization or spatial distribution of mucosal immune activity. This interpretation remains associative, but it is consistent with the broader concept that ageing alters mucosal immune structure and function [10, 39, 40]. Overall, our findings support the emerging view that ageing is spatially heterogeneous, with localized histological microenvironments exhibiting disproportionate ageing-associated remodeling.

### Cell-level features refine model attention

The improvement obtained by combining ABMIL attention with ClassPose-derived features suggests that cellular information provides complementary signal beyond mesoscopic image embeddings. Foundation-model features capture regional tissue architecture, whereas ClassPose explicitly encodes cell-type composition and morphology. By adding this information at the patch level, the fused model appears better able to distinguish visually similar regions that differ in cellular state.

Consistent with this interpretation, the fused model assigned a larger fraction of total attention to top-ranked patches. This suggests that cell-level features help refine the spatial allocation of attention toward more informative tissue regions. These findings do not imply that attention is a causal explanation, but they support the use of attention as a practical localization tool for identifying ageing-enriched histological microenvironments.

### Attention-guided histology as a strategy for spatial omics

A broader implication of this work is that H&E-based attention maps may help guide more expensive spatial molecular assays. Spatial transcriptomics and spatial proteomics preserve tissue architecture, but they are typically applied to selected regions rather than entire biopsy specimens [41, 42]. In cancer, region selection is often guided by obvious tumor morphology. In normal ageing tissue, however, informative regions are less visually obvious.

Our results suggest that model-attended regions could provide a data-driven strategy for selecting regions of interest for downstream molecular profiling. Rather than sampling histologically normal tissue at random, attention-guided selection could enrich spatial transcriptomic or proteomic experiments for regions most likely to contain age-associated biological variation. In this way, routine histology could serve as a practical bridge between low-cost tissue screening and high-resolution molecular characterization. With further validation, histology-derived ageing signatures may also provide scalable biomarkers of accelerated tissue ageing, mucosal immune dysfunction, or age-associated disease susceptibility using routine pathology archives.

### Model discordance may reveal tissue biological ageing phenotypes

Chronological age is an imperfect proxy for biological ageing. In this context, model errors may be biologically informative rather than purely technical. A sample from a younger individual predicted as older may reflect an older-appearing tissue phenotype, whereas an older individual predicted as younger may retain features of preserved tissue architecture.

Such discordant cases could be used to stratify individuals with potentially accelerated or decelerated tissue ageing, analogous to age-acceleration concepts used in molecular ageing clocks [4, 5, 43]. Future studies could focus on these false-positive and false-negative cases to determine whether they correspond to distinct molecular, inflammatory, or clinical phenotypes.

### Future directions

Our results show that routine H&E slides can reveal interpretable ageing-associated tissue patterns before molecular profiling is applied. This supports a histology-first strategy in which large pathology archives are used to screen for ageing-related phenotypes and nominate tissue compartments for deeper study.

The compartment-specific signals identified here generate focused hypotheses for follow-up. Future studies integrating histology-derived ageing signatures with spatial transcriptomics, methylation clocks, proteomics, and clinical outcomes may enable multi-modal characterization of tissue biological age. Importantly, these assays could be directed to model-attended regions rather than applied broadly or randomly across histologically normal tissue.

More broadly, this framework may help bridge routine histology and molecular ageing research by using inexpensive H&E images to localize biologically informative regions, prioritize selected samples, and guide higher-resolution experimental characterization.

## 4 Methods

### Cohort assembly and temporal data split

The study cohort consisted of normal colon biopsy WSIs collected between 2022 and 2025. Formalin-fixed, paraffin-embedded (FFPE) tissue blocks were sectioned and stained with hematoxylin and eosin (H&E) following standard protocols, and WSIs were acquired at a spatial resolution of 0.25 *µ*m per pixel using a Philips Ultra Fast Scanner (UFS). The study included normal colon tissue from the large intestine, including cases with clinical background or suspicion of inflammatory bowel disease, Crohn’s disease, ulcerative colitis, colitis, or macroscopic polyps, provided that the biopsy was histologically normal. Cases were excluded if they contained tissue adjacent to adenocarcinoma, diverticulitis-associated tissue, an anastomotic ring, or colostomy tissue; slides were additionally omitted if they were blurry, lacked tissue, or contained non-colon tissue. After applying these criteria, the final dataset comprised 631 patients and 848 WSIs. A temporal split was used for model development and evaluation: slides from 2022-2024 were assigned to the training set, comprising 536 patients and 713 WSIs, while slides from 2025 were held out as an independent test set, comprising 95 patients and 135 WSIs.

### Whole-slide preprocessing and patch embedding

Whole-slide image (WSI) preprocessing was performed using the TRIDENT pipeline, a scalable framework for computational pathology workflows that supports tissue segmentation, patch coordinate generation, and patch-level feature extraction using pathology foundation models [44–46]. Using TRIDENT, each WSI was first processed to identify tissue-containing regions and exclude background. Tissue regions were then tessellated into non-overlapping image patches, with the magnification, patch size, and encoder selected according to the downstream analysis.

For attention-based MIL modeling, WSIs were tessellated at the magnification used by the corresponding foundation-model encoder, and patch-level embeddings were extracted using the selected pretrained pathology foundation model. Patch coordinates were stored in the level-0 WSI coordinate system 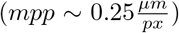. This coordinate-based representation allowed analyses performed at different magnifications to be mapped into a common slide reference frame.

Before patch embedding, patch appearance was normalized to reduce stain variability across slides. We used a multi-target Macenko stain-normalization strategy, which extends the original Macenko normalization approach by using multiple reference targets rather than a single representative target image [47, 48]. Reference targets were selected from the training set only to avoid information leakage. Specifically, we sampled 200 training slides and selected 5 patches from each slide, yielding a target bank of 1,000 reference patches. Multi-target normalization was applied in average-post-normalization mode, in which normalized patch appearances obtained relative to multiple targets are averaged after normalization. The normalized patches were then passed to the corresponding foundation model for feature extraction.

This preprocessing scheme was applied consistently within each experimental setting, while allowing the magnification, patch size, and foundation model to vary according to the specific analysis.

### Attention-based multiple instance learning (ABMIL)

We employed an attention-based multiple instance learning (ABMIL) framework to perform slide-level prediction from the patch-level embeddings extracted as described above [20]. Each WSI was represented as a bag of patch embeddings 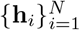, where *N* denotes the number of extracted tissue patches.

Patch-level features were obtained using a pretrained foundation model, producing a *D*-dimensional embedding for each patch. These embeddings were aggregated using a learnable attention mechanism:

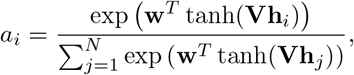

where *a*_*i*_ denotes the normalized attention weight assigned to patch *i*, and **V** and **w** are learnable parameters.

The slide-level representation was computed as a weighted sum:

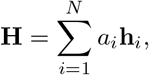

which was subsequently passed through a fully connected classifier to predict the target label. Model parameters were optimized using cross-entropy loss with class-balanced weighting to account for imbalanced age distributions.

We trained models using a cross-validation framework with out-of-fold evaluation to obtain unbiased performance estimates. Performance was assessed using area under the receiver operating characteristic curve (AUC), average precision (AP) and F_1_ score.

### PANTHER-based tissue segmentation

PANTHER is a prototype-based representation framework that partitions histological patches into a discrete set of recurrent tissue phenotypes based on similarity in feature space [26].

Prototype construction was performed using Virchow2 2560-dimensional patch embeddings extracted from 224 ×224 pixel patches at 40× magnification. Prototypes were learned using the PANTHER K-means prototyping mode on the train-set only. We specified *C* = 7 prototypes and sampled 100,000 patch embeddings for prototype construction algorithm. This produced seven data-driven morphological prototypes representing recurring tissue patterns in the training cohort.

After prototype construction, each 40× patch was assigned a PANTHER label according to the prototype with the highest assignment probability, i.e., by taking the arg max over the prototype probability vector for that patch. The resulting labels were therefore treated as categorical tissue-compartment annotations. The seven prototypes were subsequently reviewed and named by pathologists according to their dominant histological appearance, as shown in Fig. 4c: Empty Patch, Connective Tissue, Colon Crypt Epithelium, Crypt-Connective Overlap, Lymphoid Follicle, Tissue Edge, and Sub-Mucosal Tissue.

This mapping enabled aggregation of patch-level features and model attention scores within consistent histological compartments. Prototype labels were used to stratify both attention distributions and cellular feature analyses, allowing downstream statistical comparisons to be performed within PANTHER-defined compartments and thereby reducing the effect of tissue heterogeneity.

### Cell-level feature extraction using ClassPose

Cellular features were extracted from WSIs using ClassPose, a deep learning-based framework for cell detection, segmentation, and classification in histopathology images [27]. ClassPose was applied to each WSI using tiled inference with a tile size of 1024 pixels, an overlap of 64 pixels and a batch size of 8 and. The final ClassPose output included deduplicated cell detections, such that each detected cell was represented once in the exported results. For each cell, ClassPose provided a predicted cell-type label, centroid coordinates, contour geometry, and morphological measurements.

ClassPose outputs were exported as GeoJSON files containing cell centroids and cell contours. The centroid GeoJSON was used as the primary source for cell position, cell type, area, and perimeter. Cell positions were obtained from the point geometry, with centroid coordinates from the measurement fields used as fallback values when needed. Cell area and perimeter were parsed directly from the per-cell measurement fields. Cell-type labels were extracted from the classification field and mapped to six predefined cellular classes: epithelial cells, lymphocytes, neutrophils, plasma cells, eosinophils, and connective (stromal) cells.

To integrate cellular information with the patch-based MIL framework, ClassPose detections were aggregated at the level of the foundation model (FM) image patches. FM features were extracted from non-overlapping patches. Since ClassPose coordinates, as FM tessellation, were reported in level-0 coordinates, each cell was assigned to a FM patch according to whether its centroid fell within the corresponding level-0 patch boundaries. Because the FM patches were non-overlapping, each cell contributed to at most one patch.

For each patch, cell-derived features were computed separately for each cell type and for all cells combined (as a “Total” cell type). For a given patch and cell group *g*, where *g* denotes either one of the six cell types or the aggregate set of all cells, we computed a 9-dimensional feature vector:

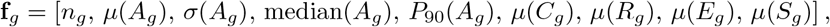

where *n*_*g*_ is the number of cells in group *g, A*_*g*_ denotes the set of cell areas, *µ*(·) denotes the mean, *σ*(·) denotes the standard deviation, and *P*_90_(·) denotes the 90th percentile.

Shape descriptors were computed at the single-cell level and then averaged within each patch and cell group. Circularity was defined as

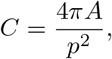

where *A* and *p* denote the cell area and perimeter, respectively. Shape irregularity was quantified using the perimeter-to-area ratio

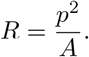

Elongation was estimated from the contour geometry by performing principal component analysis on the polygon vertices and computing

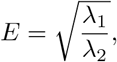

where *λ*_1_ ≥ *λ*_2_ are the eigenvalues of the covariance matrix of the contour coordinates. Solidity was computed as

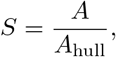

where *A*_hull_ is the area of the convex hull of the cell contour.

The final ClassPose-derived patch representation consisted of 9 features for each of the six cell types and an additional 9 features computed across all detected cells, resulting in a 63-dimensional cellular feature vector per patch. Count and area-derived features were transformed using a log(1+*x*) transformation to reduce skewness and stabilize feature magnitudes. The resulting cellular feature vector was concatenated with the corresponding UNI patch embedding to form the final patch-level representation:

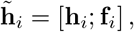

where **h**_*i*_ is the FM image embedding for patch *i*, **f**_*i*_ is the corresponding ClassPose-derived cellular feature vector, and [·;·] denotes feature concatenation. These fused patch embeddings were used as input to the downstream ABMIL model.

### Mapping attention scores to PANTHER-defined tissue compartments

Attention scores were generated at the 20× patch level by the pathological foundation model (FM), whereas PANTHER prototype labels were assigned to 40× patches. We therefore mapped attention scores from the 20× grid to the 40× PANTHER grid using spatial overlap in the common WSI coordinate system.

For slide *s*, let

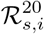

denote the spatial rectangle of the *i*-th 20× attention patch, and let

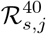

denote the spatial rectangle of the *j*-th 40× PANTHER patch. The corresponding ABMIL attention score for the 20 × patch is denoted by *a*_*s*,*i*_. When attention logits were used, they were converted to normalized attention weights by applying a slide-level softmax:

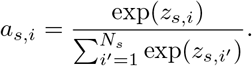

Because 20 ×and 40× patches have different physical sizes, each 40× PANTHER patch may overlap one or more 20× attention patches. For each 40× patch, we computed its mapped attention score as the area-weighted average of the overlapping 20× attention scores:

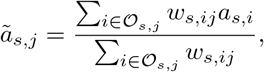

Where

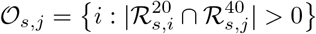

is the set of 20× patches overlapping the 40× patch, and

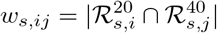

is the overlap area. Patches with no valid overlap were excluded from downstream attention-compartment analyses.

The physical scale difference between patch grids was accounted for using the known microns-per-pixel values of each magnification. In this study, 40× patches were extracted at approximately 0.25 *µ*m per pixel, whereas 20× patches corresponded to approximately 0.50 *µ*m per pixel. Patch coordinates were therefore transformed to a common coordinate system before calculating overlap areas.

Each 40× patch was associated with a PANTHER prototype label

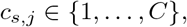

where *C* = 7 in the main PANTHER analyses. For each slide and prototype label *c*, we defined the set of valid 40× patches assigned to that prototype as

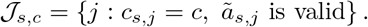

The mean mapped attention score for prototype *c* in slide *s* was then computed as

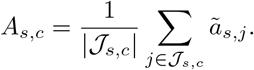

These per-slide prototype attention scores were used to quantify whether model attention was preferentially allocated to specific PANTHER-defined histological compartments.

In parallel, prototype proportions were computed as

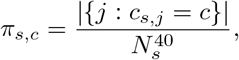

where 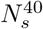 is the total number of 40× PANTHER patches in slide *s*. These proportions were used to compare mesoscopic tissue composition between age groups.

### Definition of high- and low-attention patches within PANTHER compartments

To test whether age-associated cellular features were enriched in regions prioritized by the model, we selected high- and low-attention patches within each slide and PANTHER compartment. Importantly, this selection was performed separately for each slide and each prototype label, rather than globally across the dataset, to avoid confounding by slide size or prototype abundance.

For each slide *s* and PANTHER prototype *c*, 40× patches in *J*_*s*,*c*_ were ranked by their mapped attention scores *ã*_*s*,*j*_. For a given fraction *p*, the number of selected patches was defined as

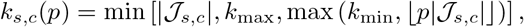

where *k*_min_ specifies the minimum number of patches selected per slide-prototype pair and *k*_max_ is an optional upper bound used to prevent large compartments from dominating the analysis.

The high-attention set was defined as the *k*_*s*,*c*_(*p*) patches with the largest mapped attention scores:

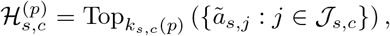

and the low-attention set was defined analogously as the *k*_*s*,*c*_(*p*) patches with the smallest mapped attention scores:

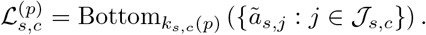

The main analysis used *p* = 0.05, corresponding to the top and bottom 5% of patches within each slide-prototype pair. Sensitivity analyses were performed using *p* = 0.01 and *p* = 0.10. The same value of *p* was used for high- and low-attention selections.

For each ClassPose-derived feature *f*, we then computed one slide-level summary value within each prototype and attention stratum:

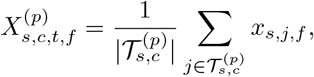

Where

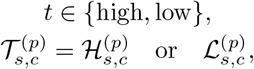

and *x*_*s*,*j*,*f*_ denotes the ClassPose-derived feature value for 40× patch *j*.

Statistical comparisons between young and old groups were then performed across slides for each feature-prototype-attention stratum combination. This ensured that the unit of statistical analysis was the slide, rather than individual patches, and avoided pseudo-replication from multiple patches within the same WSI.

## Extended Data

**Supplementary Fig. 1.**
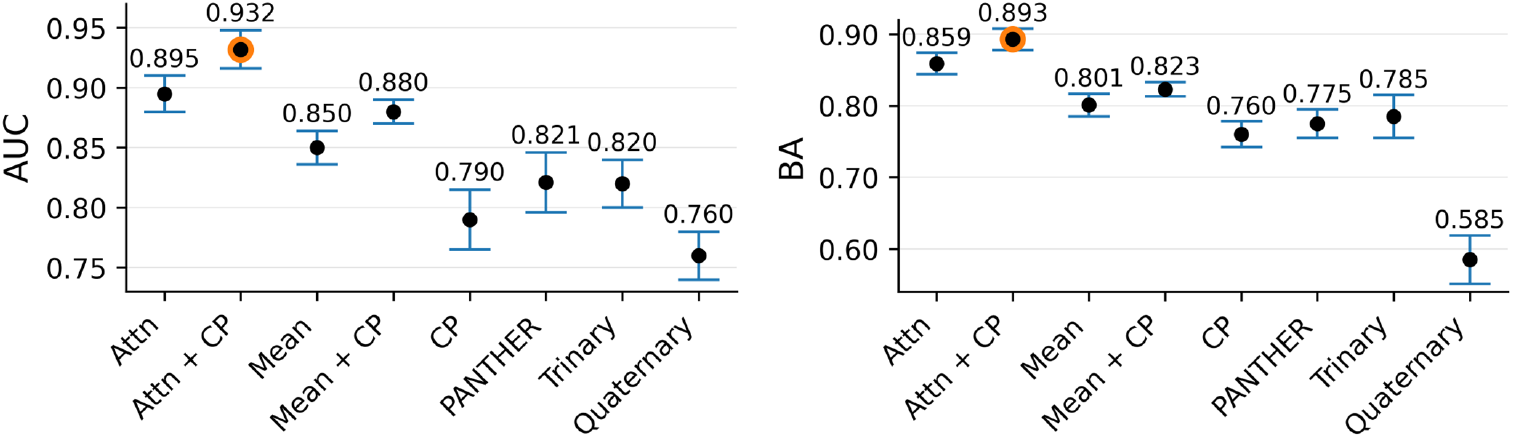
Performance across repeated training runs. Same analysis as Fig. 3a,b, with markers denoting the mean over *N* = 30 runs with different random initializations and error bars indicating the standard deviation, demonstrating robustness to training initialization.

**Supplementary Fig. 2.**
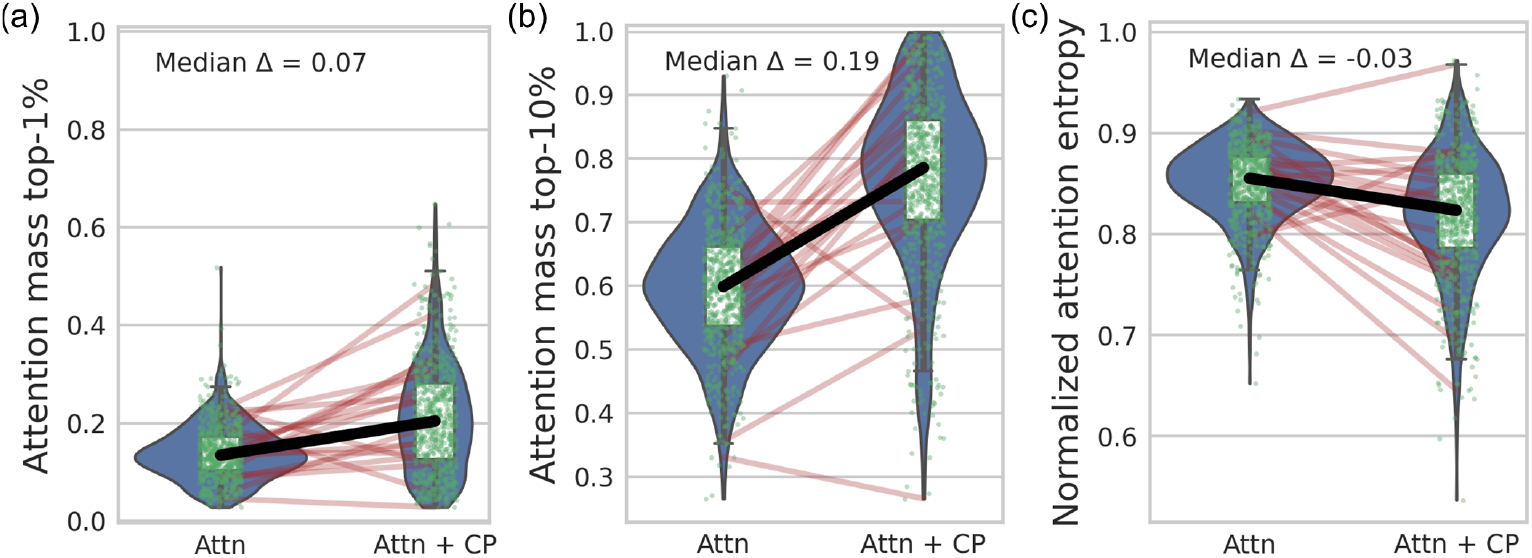
Increased attention concentration and reduced entropy after feature fusion. (a,b) Same analysis as Fig. 5d. Fraction of total attention mass assigned to the top 1% (a) and top 10% (b) highest-attention patches per slide, comparing the attention-only (Attn) and fused (Attn + CP) models. Each point represents a slide; lines connect paired observations. The fused model consistently allocates a larger share of attention to top-ranked patches, indicating increased concentration of decision weight. Median differences (Δ) are indicated. (c) Normalized attention entropy per slide, showing reduced entropy in the fused model, consistent with a more peaked attention distribution. Together, these results demonstrate that integrating cell-level features sharpens spatial attention and concentrates model focus on a smaller subset of histological regions.

## References

[1] López-Otín, C., Blasco, M. A., Partridge, L., Serrano, M. & Kroemer, G. The hallmarks of aging. Cell 153, 1194–1217 (2013).

[2] López-Otín, C., Blasco, M. A., Partridge, L., Serrano, M. & Kroemer, G. Hallmarks of aging: An expanding universe. Cell 186, 243–278 (2023).

[3] Levine, M. E. et al. An epigenetic biomarker of aging for lifespan and healthspan. Aging (albany NY) 10, 573 (2018).

[4] Horvath, S. Dna methylation age of human tissues and cell types. Genome biology 14, 3156 (2013).

[5] Lu, A. T. et al. Dna methylation grimage strongly predicts lifespan and healthspan. Aging (albany NY) 11, 303 (2019).

[6] Funk, M. C., Zhou, J. & Boutros, M. Ageing, metabolism and the intestine. The EMBO Reports 21, EMBR202050047 (2020).

[7] Choi, J. & Augenlicht, L. H. Intestinal stem cells: guardians of homeostasis in health and aging amid environmental challenges. Experimental & Molecular Medicine 56, 495–500 (2024).

[8] Baidoo, N. & Sanger, G. J. The ageing of the human lower bowel. British Journal of Hospital Medicine 86, 1–16 (2025).

[9] Branca, J. J., Gulisano, M. & Nicoletti, C. Intestinal epithelial barrier functions in ageing. Ageing research reviews 54, 100938 (2019).

[10] Zheng, H. et al. The impact of aging on intestinal mucosal immune function and clinical applications. Frontiers in immunology 13, 1029948 (2022).

[11] Franceschi, C., Garagnani, P., Parini, P., Giuliani, C. & Santoro, A. Inflammaging: a new immune–metabolic viewpoint for age-related diseases. Nature Reviews Endocrinology 14, 576–590 (2018).

[12] Wiley, C. D. & Campisi, J. The metabolic roots of senescence: mechanisms and opportunities for intervention. Nature metabolism 3, 1290–1301 (2021).

[13] Daly, A. C. et al. Tissue and cellular spatiotemporal dynamics in colon aging. Nature biotechnology 1–13 (2025).

[14] Chen, R. J. et al. Towards a general-purpose foundation model for computational pathology. Nature Medicine (2024).

[15] Vorontsov, E. et al. A foundation model for clinical-grade computational pathology and rare cancers detection. Nature medicine 30, 2924–2935 (2024).

[16] Zimmermann, E. et al. Virchow2: Scaling self-supervised mixed magnification models in pathology. arXiv preprint arXiv:2408.00738 (2024).

[17] Xu, H. et al. A whole-slide foundation model for digital pathology from real-world data. Nature 630, 181–188 (2024).

[18] Meng, R. et al. Machine-learning models based on histological images from healthy donors identify imageqtls and predict chronological age. Proceedings of the National Academy of Sciences 122, e2423469122 (2025).

[19] Peng, Z. et al. Benchmarking pathology foundation models for non-neoplastic pathology in the placenta. medRxiv (2025).

[20] Ilse, M., Tomczak, J. M. & Welling, M. Attention-based deep multiple instance learning (2018).

[21] Lu, M. Y. et al. Data-efficient and weakly supervised computational pathology on whole-slide images. Nature biomedical engineering 5, 555–570 (2021).

[22] Chen, R. J. et al. Scaling vision transformers to gigapixel images via hierarchical self-supervised learning, 16144–16155 (2022).

[23] Song, A. H. et al. Artificial intelligence for digital and computational pathology. Nature Reviews Bioengineering 1, 930–949 (2023).

[24] Lu, M. Y. et al. A visual-language foundation model for computational pathology. Nature medicine 30, 863–874 (2024).

[25] Ding, T. et al. A multimodal whole-slide foundation model for pathology. Nature medicine 1–13 (2025).

[26] Song, A. H. et al. Morphological prototyping for unsupervised slide representation learning in computational pathology, 11566–11578 (2024).

[27] Mandal, S., de Almeida, J. G., Papanikolaou, N. & Graham, T. A. Classpose: foundation model-driven whole slide image-scale cell phenotyping in h&e. bioRxiv 2025–12 (2025).

[28] Rudin, C. Stop explaining black box machine learning models for high stakes decisions and use interpretable models instead. Nature machine intelligence 1, 206–215 (2019).

[29] Band, S. S. et al. Application of explainable artificial intelligence in medical health: A systematic review of interpretability methods. Informatics in Medicine Unlocked 40, 101286 (2023).

[30] Franceschi, C. & Campisi, J. Chronic inflammation (inflammaging) and its potential contribution to age-associated diseases. Journals of Gerontology Series A: Biomedical Sciences and Medical Sciences 69, S4–S9 (2014).

[31] Straub, R. H. & Schradin, C. Chronic inflammatory systemic diseases: An evolutionary trade-off between acutely beneficial but chronically harmful programs. Evolution, medicine, and public health 2016, 37–51 (2016).

[32] Sanada, F. et al. Source of chronic inflammation in aging. Frontiers in cardiovascular medicine 5, 12 (2018).

[33] Ferrucci, L. & Fabbri, E. Inflammageing: chronic inflammation in ageing, cardiovascular disease, and frailty. Nature Reviews Cardiology 15, 505–522 (2018).

[34] Tran, L. & Greenwood-Van Meerveld, B. Age-associated remodeling of the intestinal epithelial barrier. Journals of Gerontology Series A: Biomedical Sciences and Medical Sciences 68, 1045–1056 (2013).

[35] Baidoo, N. & Sanger, G. J. Age-related decline in goblet cell numbers and mucin content of the human colon: implications for lower bowel functions in the elderly. Experimental and Molecular Pathology 139, 104923 (2024).

[36] Baidoo, N., De Rasis, E., Tahiri, D. & Sanger, G. J. The impact of ageing on the structural compositions of the mucosa of human colon. Tissue and Cell 97, 103090 (2025).

[37] Fenton, T. M. et al. Immune profiling of human gut-associated lymphoid tissue identifies a role for isolated lymphoid follicles in priming of region-specific immunity. Immunity 52, 557–570 (2020).

[38] Mörbe, U. M. et al. Human gut-associated lymphoid tissues (galt); diversity, structure, and function. Mucosal immunology 14, 793–802 (2021).

[39] Kato, H. et al. Lack of oral tolerance in aging is due to sequential loss of peyer’s patch cell interactions. International immunology 15, 145–158 (2003).

[40] Donaldson, D. S., Shih, B. B. & Mabbott, N. A. Aging-related impairments to m cells in peyer’s patches coincide with disturbances to paneth cells. Frontiers in Immunology 12, 761949 (2021).

[41] Jain, S. & Eadon, M. T. Spatial transcriptomics in health and disease. Nature reviews nephrology 20, 659–671 (2024).

[42] Wu, M. et al. Spatial proteomics: unveiling the multidimensional landscape of protein localization in human diseases. Proteome science 22, 7 (2024).

[43] Lu, A. T. et al. Dna methylation grimage version 2. Aging (Albany NY) 14, 9484 (2022).

[44] Mahmood Lab. TRIDENT: Toolkit for large-scale whole-slide image processing. https://github.com/mahmoodlab/TRIDENT (2025). Accessed 2026-05-05.

[45] Zhang, A., Jaume, G., Vaidya, A., Ding, T. & Mahmood, F. Accelerating data processing and benchmarking of ai models for pathology. arXiv preprint arXiv:2502.06750 (2025).

[46] Vaidya, A. et al. Molecular-driven foundation model for oncologic pathology. arXiv preprint arXiv:2501.16652 (2025).

[47] Macenko, M. et al. A method for normalizing histology slides for quantitative analysis, 1107–1110 (IEEE, 2009).

[48] Ivanov, D., Barbano, C. A. & Grangetto, M. Multi-target stain normalization for histology slides, 36–44 (Springer, 2024).

